# Boredom begets creativity: a solution to the exploitation-exploration trade-off in predictive coding

**DOI:** 10.1101/104521

**Authors:** Jaime Gomez-Ramirez, Tommaso Costa

## Abstract

Here, we investigate whether systems that minimize prediction error e.g. predictive coding, can also show creativity, or on the contrary, prediction error minimization unqualifies for the design of systems that respond in creative ways to non recurrent problems. We argue that there is a key ingredient that has been overlooked by researchers that needs to be incorporated to understand intelligent behavior in biological and technical systems. This ingredient is boredom. We propose a mathematical model based on the Black-Scholes-Merton equation which provides mechanistic insights into the interplay between boredom and prediction pleasure as the key drivers of behavior.

## 1 Introduction

The value in building artificial systems with optimal predictive power is beyond question. Robots in real world missions without the capacity to infer the state of the world are unreliable and doomed to a short existence. In biological systems, the idea that organisms organize sensory data into an internal model of the outside world, goes back to the early days of experimental psychology. In Helmholtz’s *Handbook of Physiological Optics* published in 1866, it is argued that the brain unconsciously adjusts itself to produce a coherent experience. According to this view, our perceptions of external objects are images or better said, symbols, that do not resemble the referenced objects. Helmoltz’s view of perception as a process of probabilistic inference, in which sensory causes need to be inferred based upon changes of body states, has become a major tenet in a number of disciplines, including computational neuroscience (Dayan and Abbott, 2002), cybernetics (Ashby, 2015), cognitive psychology (Neisser, 2014) and machine learning (Neal and Hinton, 1998).

A recent incarnation of this theory of perception is the Helmholtz’s machine postulated by Dayan, Hinton and Zemel (Dayan et al., 1995), (Dayan and Hinton, 1996). The brain is here conceptualized as a statistical inference engine whose function is to infer the causes of sensory input. Under this scheme, the workings of the brain encode Bayesian principles. Due in part to the ever increasing computational power of computers, Bayesian approaches alike to the Helmholtz’s machine have become the workhorse for studying how the nervous system operates in situations of uncertainty (Rao and Ballard, 1999), (Knill and Pouget, 2004), (Friston, 2012). The main rationale is that the nervous system maintains internal probabilistic models which are continuously updated in the light of their performance in predicting the upcoming suite of cues.

Predictive coding is a form of differential coding where the signal of interest is the difference between the actual signal and its prediction. This technique exploits the fact that under stationary and ergodic assumptions^1^, the value of one data point e.g., a pixel, regularly predicts the value of its nearest neighbors. Accordingly, the variance of the difference signal is reduced compared to the original signal, making differential coding an efficient way to compress information (Shi and Sun, 1999). In a general sense, predictive coding is a Bayesian approach to brain function in which the brain is conceived as a device trained to do error correction. Predictive coding aims at reducing redundancy for signal transmission efficiency and it is been proposed as a unifying mathematical framework for understanding information processing in the nervous system (Friston, 2010), (Huang and Rao, 2011). Predictive coding has been used to model spatial redundancy in the visual system (Srinivasan et al., 1982), temporal redundancy in the auditory system (Baldeweg, 2006) and the mirror neuron system (Kilner et al., 2007).

Predictive coding is a “neuronally plausible implementation scheme” (Schwartenbeck et al., 2013) of the free energy minimization principle which is a theoretical formulation that in essence states that biological systems always behave under the imperative of minimizing surprise. In a series of articles spanning over one decade, Friston and collaborators have proposed a free energy principle as an unified account of brain function and behavior. The free energy minimization principle buttresses Helmholtz’s theory perception using modern-day statistical theories, namely, Bayesian filtering (Friston, 2005), Maximum entropy principle (Jaynes, 2003) and variational free energy (Hinton and van Camp, 1993) (Table 1).

**Table I.** Definition of concepts.

|  |  |
| --- | --- |
| Helmoltz machine | A neural network that represents its inputs with a minimal-length description using recognition connections, which run from inputs to outputs, and generative connections, which run from outputs back to inputs. Helmholtz machines infer the cause of its input using variational free energy as a proxy of surprise |
| Bayesian brain | The brain tries to infer the causes of our sensations based on a generative model of the world. The Bayesian brain is a corollary of the free-energy principle |
| Predictive coding | A Bayesian approach to brain function in which the brain is conceived as a device trained to do error correction. It is a brain inspired implementation of the free energy minimization principle |
| Free energy minimization principle | Mathematical framework of Helmholtz theory of perception and action. The main assumption is that adaptive systems resist a natural tendency to disorder by minimizing the free energy |

The actual relevance and soundness of the free energy principle to explain decision making in organisms is being contested. Critics argue that if biological systems behave in the way that free energy minimization prescribes-minimizing surprises over the states visited-they will inevitably seek the most predictable habitat, for example, a corner in a dark room, and they will stay there ad infinitum. This mental experiment is being called the "dark room problem" (Friston et al., 2012) and shows that the imperative of organisms to minimize surprise, put forward by the free energy minimization principle, is at odds with easily recognizable features of organisms such as exploration or creativity.

Friston’s way out of the "dark-room problem" is as follows, probabilities are always conditional to the system’s prior information, thus, a system equipped with a generative model (priors and a likelihood function) that dislikes dark rooms rather than being stuck in a corner minimizing its prediction error, will walk away in order to sample the external world according to its own priors. Crucially, surprise or surprisal ^2^over states (*S*) is always conditional to a given specific generative model (*m*). The surprise over *S* is always conditional to the model *m*, *H*(*S*|*m*), which is obviously different to the marginal surprise over the states, *H*(*S*). Different systems acting in identical environments might disagree in what is surprising and what is not according to their priors. Therefore, organisms with a tendency to explore and take risks will not necessarily purse a bland and uneventful existence as the "dark-room problem" suggests.

But where the priors come from and how they are shaped by the environment is never said in the predictive coding framework. This is indeed the crux of the matter in Bayesian statistics. The translation of subjective prior beliefs into mathematically formulated prior distributions is an ill-defined problem (Gomez-Ramirez and Sanz, 2013). And yet, the minimization of surprise is a sufficient condition for keeping the system within an admissible set of states. A bacterium, a cockroach, a bird and a human being all have in common that in order to persevere in their actual forms, they must limit their physiological states, that is, organisms constrain their phenotype in order to resist disorder. Homeostasis is the control mechanism in charge of keeping the organism’s internal conditions stable and within bounds. Survival depends on the organism’s capacity to maintain its physiology within an optimal homeostatic range (Damasio and Carvalho, 2013).

But where the priors come from and how they are shaped by the environment is never said in the predictive coding framework. This is indeed the ^2^ See the S1 Appendix for the technical definition of surprisal and implications within the free energy principle and the predictive coding framework. crux of the matter in Bayesian statistics. The translation of subjective prior beliefs into mathematically formulated prior distributions is an ill-defined problem (Gomez-Ramirez and Sanz, 2013). And yet, the minimization of surprise is a sufficient condition for keeping the system within an admissible set of states. A bacterium, a cockroach, a bird and a human being all have in common that in order to persevere in their actual forms, they must limit their physiological states, that is, organisms constrain their phenotype in order to resist disorder. Homeostasis is the control mechanism in charge of keeping the organism’s internal conditions stable and within bounds. Survival depends on the organism’s capacity to maintain its physiology within an optimal homeostatic range (Damasio and Carvalho, 2013).

This is the conundrum that this paper addresses. On the one hand, free energy minimization is conducive to achieving the homeostatic balance necessary for the organism’s survival and well-being and on the other hand, surprise minimization can not possibly be the unique modus-operandi of biological systems. Organisms that minimize the entropy of the sensory states they sample would never engage in exploration, risk-taking or creativity, for the simple reason that these behaviors might increase the prediction error. In consequence, surprise ^3^ can not be used as the unique necessary factor to explain choices under uncertainty conditions.

Here we argue that the actual quantity that is maximized is the difference between prediction error and boredom. The crucial intuition behind our model is strikingly simple. A system that minimizes prediction error is not only attentive to homeostasis and the vital maintenance functions of the body, but it also maximizes pleasure. For example, the reward effect in the appreciation of aesthetic work might come from the transition from a state of uncertainty to a state of increased predictability (Cruys and Wagemans, 2011). However, this is until the signal error becomes stationary, or in the art work example, the art work has not anymore the potential of surprising us, in that case boredom kicks in, reducing the overall value of the subjective experience. Boredom is an aversive (negative valence) emotion (Goetz et al., 2013), (Joffily and Coricelli, 2013). Thus, boredom creates the conditions to start exploring new hypothesis by sampling the environment in new and creative ways, or put in other words, boredom begets creativity. Until very recently, the function of boredom has been considered of little or no interest for understanding human functioning. This situation is rapidly changing, recent studies in human psychology shows that the experience of boredom might be accompanied by stress and increases levels of arousal to ready the person for alternatives (Posner et al., 2009) (Bench and Lench, 2013).

The rest of the paper is structured as follows. Section 2 introduces a mathematical model that extends and complements predictive coding. Surprise minimization in any of its equivalent forms such as free energy minimization and marginal likelihood maximization is not a sufficient but a necessary *explanans* of biological behavior. Section 3 presents the simulations of the model to help have an intuitive grasping of the mathematical model based on the Black-Scholes-Merton equation. This statistical model, though conceived for a very different problem (financial options pricing) allows us to elegantly capture the interplay between prediction and boredom. In section 4 we discuss the limitations of free energy minimization and its neuronal implementation, prediction coding, in relation with the previous results.

## 2 Methods

In this section we build a mathematical model to explain intelligent behavior as the maximization of the subjective experience. The subjective experience consists of two terms with opposed valence, prediction pleasure and boredom. Prediction pleasure is a positive or hedonic state and boredom is a negative emotional state. In short, what organisms do is to maximize subjective experience, and in order to achieve that objective they tend to minimize surprise as predictive coding correctly claims, while at the same time diminishing boredom, a negative emotion that arises during monotonous tasks or in environments with low entropy. The rationale behind this is that organisms maximize subjective experience by making prediction pleasure as large as possible while keeping boredom low ^4^. In this view, organisms do not exclusively operate in prediction mode, sooner or later, depending on the intrinsic agent’s motivations and how they match with the environment, the marginal utility of prediction will decrease and the organism will switch to exploration mode, that is, the organism will become less concerned with predicting its current state, and will be prone to visit surprising states that overall increase its well being. In line with this idea, Schmidhuber (Schmidhuber, 2010) has proposed a general formal theory of fun and creativity based on the discovery of novel or surprising patterns which according to the model, maximizes intrinsic reward allowing for improved prediction. Kakade and Dayan (Kakade and Dayan, 2002) have suggested that dopamine neurons encode "reward bonuses", playing a fundamental role in explorative behavior.

We start by defining the utility function ^5^ that agents maximize as

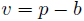

where *v* is the subjective experience and *p* and *b* represent prediction pleasure and boredom, respectively. It seems clear from equation 1 that the larger the prediction pleasure (*p*) the greater the value of the subjective experience (*v*), limited by the boredom (*b*) that prediction brings in. When the prediction pleasure is greater than the boredom the subjective experience is overall positive or pleasant, on the contrary, when the boredom exceeds the prediction pleasure, experience is negative or “painful”.

We need now to be more precise in the formulation of the terms included in equation 1. Reinforcement learning is the problem faced by an agent that must learn to predict the value of future events through the computation of the difference between one’s rational expectations of future rewards and any information that leads to a revision of expectations (Glimcher, 2011). Prediction error is a function of prediction error and time, specifically prediction pleasure is the inverse of prediction error. Thus, the instantaneous subjective experience *v*_*t*_ is calculated as the difference between the instantaneous pleasure *p*_*t*_ and the boredom *b*_*t*_, which in our model is assumed to be constant or *b_t_* = *k*. The boredom constant *k* represents the agent’s disposition to get bored and is therefore an inherent property of the system or *causa sui*. Prediction pleasure, on the other hand, is directly calculated from the prediction error. Prediction pleasure at time *t, p*_*t*_, is the reciprocal of prediction error at time *t*, *ϵ*_*t*_, that is, 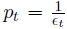. Accordingly, the value of the experience at time *t* is the difference between the prediction pleasure at *t* minus the boredom component, that is,

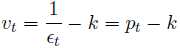

A reasonable assumption is that the prediction error describes a generalized Wiener process (Ross, 1996). A Wiener process is a particular type of Markov process which is a stochastic process where only the current value of a random variable is relevant for future prediction. Thus, we define the prediction error as *ϵ* a generalized Wiener process

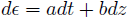

where *ϵ* is a random variable that represents the prediction error, *a* is the drift or the mean change per unit time, *b* the variance per unit time and *dz* is a Wiener process with zero drift and 1.0 variance rate. Since the drift is equal to zero, the expected value of *z* is zero, that is, at any future time, *z* is expected to be equal to its current value. The variance rate of 1.0 means that the variance of the change in *z* in a time interval of length *T* is equal to *T* i.e. the variance rate grows proportionally to the maturity time *T*.

If we additionally assume that the variability of the "return" of prediction error in a short period of time is the same regardless of the actual value of the prediction error *ϵ*, e.g. we are equally uncertain about having a gain of for example, 10% in prediction error when the prediction error is 1.6 and when it is 5.5, then the prediction error percentage change is defined as

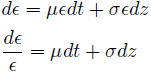

where *µ* is the expected rate of return, i.e. the percentage of change in the prediction error for one time period and *σ* is the volatility of prediction error. For example, *µ* = 0.1 means that prediction error is expected to increment by a 10%. Following the assumption that the prediction error follows a Wiener process, *µ* = 0 and *σ* = 1.0.

Remind that in equation 1 we defined the subjective experience *v* as the difference between prediction pleasure *p* and boredom *b*. The prediction pleasure *p* is a function of the underlying stochastic variable *ϵ* or prediction error. The Itô lemma allows us to characterize a function of a variable that follows a Itô process (Ito, 1951). Since the prediction error *ϵ* is a generalized Wiener process, it can be modeled as a Itô process,

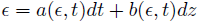

where *dz* is a Wiener process whose drift *a* and variance rate *b*, rather than constant, are functions of *ϵ* and *t*. The Itô lemma shows that a function (*f*) of a Itô process (*x*) follows as well the Itô process described in equation 6. The demonstration can be found elsewhere (Shreve, 2010).

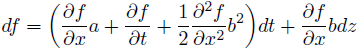

Now, substituting *f* for *p* and *x* for in equation 6 gives the prediction pleasure behavior *p* derived from the underlying prediction error. Since both *ϵ* and *p* follow geometric Brownian motion, the prediction pleasure *p* corresponds to the Itô process

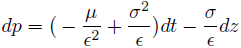

Note that equation 7 is not a generalized Wiener process because the drift rate and the variance rate are not constant. Using the Itô lemma, the process followed by *γ* = *lnϵ* when *ϵ* follows the process described in equation 5 is

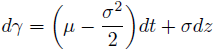

Since *µ* and *σ* are constant, *γ* = ln*ϵ* follows a general Wiener process with drift rate 
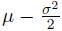
 and variance rate *σ*^2^. The change in ln between 0 and the final T normally distributed, and therefore the prediction error *ϵ* is lognormally distributed. The inverse of the prediction error or the prediction pleasure is also lognormally distributed, see the S1 Appendix for the demonstration. We use Monte-Carlo simulation to sampling random outcomes of the Itô process.

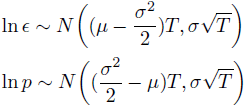

Consider now that we are interested in studying the behavior of a system with a boredom constant *k* over a time period *T*. The expected experience value at time *t* (*v*_*t*_) is its expected value at time *T* (*v*_*T*_) discounted at the rate *r*. This idea relies upon the method of asset valuation called discounted cash flow. The money in the future and now have different values because in order to correctly quantify value one needs to discount for the rate at which the money grows. For example, 100$ value asset with an annual growth rate of *r* = 10% and 5 years maturity is value today 67.3$ ^6^.

For the s, the discount factor *r* can be here understood as a prediction rate, which in essence represents how much structure there is in the outside world. For example, in an external world in which information is entirely redundant, *r* will be zero. In the other extreme of the spectrum, a fairly complex world contains a rich mosaic of patterns to be discovered by an agent equipped with the adequate perceptual, motoric and cognitive capabilities. The larger the prediction rate *r*, the more structure there is in the world. Thus, the rate *r* can be seen as a proxy for the structure of the outside world.

We define the value of the experience of an agent at time *t, t < T*, as the expected experience value defined in equation 2, discounted at rate *r*. Formally,

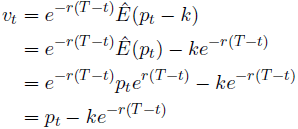

where *v*_*t*_ is the experience value at time *t*, *p*_*t*_ is the prediction pleasure at time *t*, *k* is the boredom constant and *r* is external world complexity. According to equation 10 the subjective experience at time *t, t* < *T, vt* is equal to the expected prediction pleasure minus the boredom at the final time *T* discounted at rate *r*. If the final or maturity time *T* is very far in the future, then the value of the subjective experience will be very similar to the prediction pleasure, *v*_*t,T*−*t*→∞_ = *pt* On the other hand, if the expiration date is near, the subjective experience is equal to prediction pleasure minus the boredom constant, *v*_*t,T*−*t*→0_ = *pt* − *k* Importantly, equation 10 assumes that both prediction and boredom mode are equally likely.

A more realistic model will weight the prediction and boredom terms by their respective probabilities. In order to do so we use the Black-Scholes-Merton model for option pricing. In a seemingly way as an option price is a derivative of a stock price, a subjective experience value can be calculated with the underlying prediction pleasure at a given time *t* within a time horizon *T, t < T*. We thus, borrow from the Black-Scholes-Merton model for option pricing (Black and Scholes, 1973) to model subjective experience as a "derivative".

In finance, a derivative derives its value from the performance of an underlying entity. In our model, the derivative of the subjective experience is calculated from the underlying prediction pleasure and the pain-related boredom. The Black-Scholes-Merton model will thus, help us establishing a working analytical framework to study the interplay between prediction and boredom. For a more in depth discussion on the Black-Scholes-Merton model, the reader might want to consult the S1 Appendix together with the seminal paper (Black and Scholes, 1973) and two excellent textbooks, (Hull, 2011) and (Duffie, 2001).

In the Black-Scholes-Merton option pricing model, the option is exercised only when the payoff is positive, that is to say, the stock has more value than the strike price stipulated in the contract, (*S*_*t*_−*K >* 0). In case the stock has less value than the strike price (*S*_*t*_ − *K <* 0), the holder of the option is not obligated to buy the asset. In our model, on the other hand, the subjective experience is always "exercised". This means that the experience is what it is, positive when prediction is larger than boredom and negative when the boredom exceeds the prediction pleasure. Taking this into account, we define the value of the experience as the difference between prediction and boredom, discounted and weighted by the probability of being in each mode.

Formally,

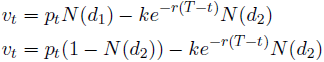

where the first term in the right side of equation 11 represents the prediction pleasure factored by the probability of being in predictive mode, *N*(*d*_1_), and the second term quantifies the pain or negative experience triggered by a boring experience in a world with a prediction rate *r* discounted at time *t* and factored by the probability of being in boredom mode, *N*(*d*_2_). The terms *N*(*d*_1_) and *N*(*d*_2_) in equation 11 are as in the Black-Scholes-Merton equation, cumulative probability distribution functions, i.e. *N*(*d*_*i*_) = *P*(*x > d*_*i*_) of the variables *d*_1_ and *d*_2_.

Assuming that the agent at any give instant can be in one of the two possible modes - prediction or boredom - we just need to define one of the two terms, for example *d*_2_, to obtain both *N*(*d*_2_) and *N*(*d*_1_) = 1 − *N*(*d*_2_)

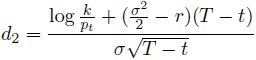

If prediction pleasure is very large compared to boredom, *p*_*t*_*/k >>* 1, *d*_2_ will be very small and *d*_1_ very large, resulting *N*(*d*_1_) → 1 and *N*(*d*_2_) → 0. In this situation, the overall experience will be positive. On the contrary, when *p*_*t*_*/k* → 0 the overall experience will be negative or dominated by boredom, i.e. *N*(*d*_2_) → 1.

## 3 Results

Equation 11 captures boredom begets creativity in the sense that boredom decreases the subjective value, possibly triggering corrective actions like exploring or wandering at the expense of reducing prediction pleasure but incrementing the subjective experience value. We run simulations for the different parameters of the model i.e. initial prediction pleasure (*p*_0_), expected rate of return 
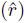
, boredom (*k*) and the drift (*µ*) and the variance per unit time (*σ*) of the prediction error. Since prediction error is assumed to follow a Wiener process, *µ* = 1 and *σ* = 0. The initial prediction pleasure, *p*_0_, and the boredom constant, *k*, can be seen as the priors. For example, all things being equal, an agent with a large ratio *k/p*_0_ is likely to have a predominantly boredom experience compared to another agent with a large *p*_0_*/k* which, on the contrary, will likely have a positive experience dominated by prediction pleasure.

The priors *p*_0_ and *k* are useful to classify agents into two categories, those with large *p*_0_*/k* enjoy predicting and can be referred to as "copiers", while those with low *p*_0_*/k* tend to get bored and fall under the category "explorers". In addition to the bias or predisposition of the agent to predict or get bored and explore given by the priors *p*_0_ and *k*, the expected rate of return 
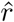
 represents the environment’s complexity. Thus, a world with large *rˆ* has more structure or patterns to be decoded by the agent than a world with low 
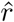
. We normalize the value of 
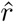
 to take values between 0 and 1. For example, for two agents, *a*_1_ and *a*_2_ in their respective environments, 
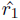
 and 
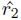
 with 
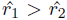
 agent *a*_1_ will need more time to get bored and eventually go to explore (*N*(*d*_2_) *> N*(*d*_1_) in equation 12) than agent *a*_2_ because the environment of *a*_1_ has more structure than the environment of *a*_2_.

Figure 1 shows a simulation of the model for a trivially predictable environment. We codify this situation with the parameter 
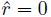
. An example of this environment is a dark room, the world here is assumed to have very low informational complexity. In this scenario, when the agent does not have any particular predisposition to predict versus to explore (*p*_0_ = *k*), prediction pleasure decays linearly and boredom remains stationary. Since the world is trivially predictable, the boredom term, which can be seen as a signal to explore, remains constant. This is because there is no structure to be discovered in the world and therefore it does not make sense to explore it (Figure 1 *a*). If the environment is trivially predictable 
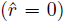
 and the agent is an explorer, that is, has bias to get bored (*p*
_0_
*/k <* 1), boredom will grow and prediction will decay, resulting in a negative experience value at the end of the period 1 *b*). Finally, when the agent is a copier, that is, has a predisposition to predict as opposed to explore (*p*_0_*/k >* 1), due to this bias, the probability of being in prediction mode is initially larger than being in boredom mode (*N*(*d*_1_) *> N*(*d*_2_)) and will continue growing to reach a maximum at the end of the period (*N*(*d*_1_) = 1*, N*(*d*_2_) = 0). The bias for predicting governs the overall experience which is always positive. (Figure 1 *c*)

Figure 2 shows simulations of the model when the world is rich on structure, that is, there are patterns to be discovered and possibly, surprising events using the definition of surprise given in the Introduction section. We codify this situation with the parameter 
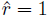
 to denote a rich world structure whose information can be compressed into meaningful patterns. When the agent does not have any particular predisposition of being in prediction or boredom mode (*p*_0_*/k* = 1), the probability of being in prediction mode, *N*(*d*_1_), is at time 0 larger than the probability of being in boredom mode, *N*(*d*_2_), because the external world is structured. At final time *T,* since the agent lacks any bias versus prediction or exploring, *N*(*d*_1_) = *N*(*d*_2_), the experience value decreases as the boredom rises and the prediction pleasure decays. The rationale behind this is that even though the agent is predicting the world and therefore having prediction pleasure, being consistently successful at predicting the world has the side effect of getting bored reducing the overall experience value (Figure 2 *a*). If the agent in an eventful world structure, 
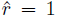
 has a predisposition to get bored (*p*_0_*/k <* 1), initially, since the world is rich in structure, the agent will be in prediction mode *N*(*d*_1_) *> N*(*d*_2_), but as the time goes on, the boredom component will exceed the prediction component and the overall experience will be negative. Thus, if the agent is an explorer in a world rich in stimuli, the experience value will become negative after some time e.g. *t* = 0.4 in figure 2 *b* and therefore it will need to take action i.e. explore the world, in order to diminish the boredom-related pain. The rationale here is that since the world has complexity (patterns to be identified by the agent), boredom will act as a signal to explore the world that will keep the agent from predicting when predicting has decreasing marginal utility. Metaphorically speaking, the agent anticipates that the "low-hanging fruit" will not last for ever, investing in new ways of reward seeking behavior(Figure 2 *b*). An agent that is a copier, (*p*_0_*/k >* 1) in an eventful world structure, 
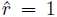
 resides in a world that suits its own personality. The subjective experience, though slowly decreasing, is always positive and importantly it will not get bored. The rationale is that the agent will content himself copying the rich structure of the world (Figure 1 *c*).

To capitulate, in both figures for either low and high world complexity, when the agent has a bias to get bored (explorer agent) the boredom component raises reducing the overall experience value (Figures 1 and 2
*b*). When the agent has a predisposition to predict (copier agent), the subjective experience increases in the dark room like scenario because the agent’s priors matches with the world easiness to predict (Figure 1 *c*), and when the informational complexity of the world is high, the overall experience will decrease but remains positive because the agent has a bias to predict and there is structure, surprises and novel patterns to be predicted in the world (Figure 2 *c*). When the agent has no priors (Figures 1 and 2 *a*), is neither a copier nor an explorer and the subjective experience decreases. In the case of a trivially predictable world (Figure 1 *a*), this is because the marginal utility of prediction pleasure will necessarily decrease. In the complex world (Figure 2 *a*), the subjective experience decreases because boredom will increase to signal that there may be more in the world that can be easily predicted.

**Figure 1:**
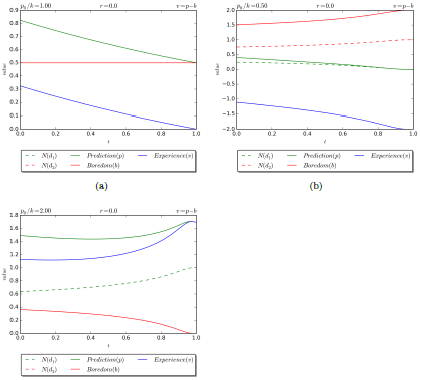
The figure shows the evolution of the probabilities of being in prediction mode, *N*(*d*_1_), boredom mode *N*(*d*_2_), the prediction pleasure (*p*), the boredom-related pain (*b*) and the experience value (*v*) for a trivial world with low complexity 
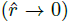
 e.g. a dark-room environment. In Figure 1-*a* there is no initial bias, (*p*_0_ = *k*), in Figure 1-*b* there is a bias that favors exploring triggered by boredom (*p*_0_*/k <* 1) and in Figure 1-*c* the bias is versus prediction against boredom (*p*_0_*/k >* 1). With the exception of 1-*c* prediction pleasure decreases driving the experience to 0 or negative values. In Figure 1-*c*, the agent is a copier in a world that is easy to copy, thus by copying the world the agent maximizes his experience. Translating these results in the dark room Gedankenexperiment, the agent will get out of the room to explore and hopefully increment the experience value that is otherwise decreasing. This occurs when either the agent has no bias (*p*_0_ = *k*, Figure 1-*a*) and when it has a bias to explore (*p*_0_*/k <* 1, Figure 1-*b*). When the agent is a copier, *p*_0_*/k >* 1, Figure 1-*c*) shows that it will stay in the dark room since it has no incentive to explore outside, boredom decreases and the overall experience driven by prediction pleasure increases.

**Figure 2:**
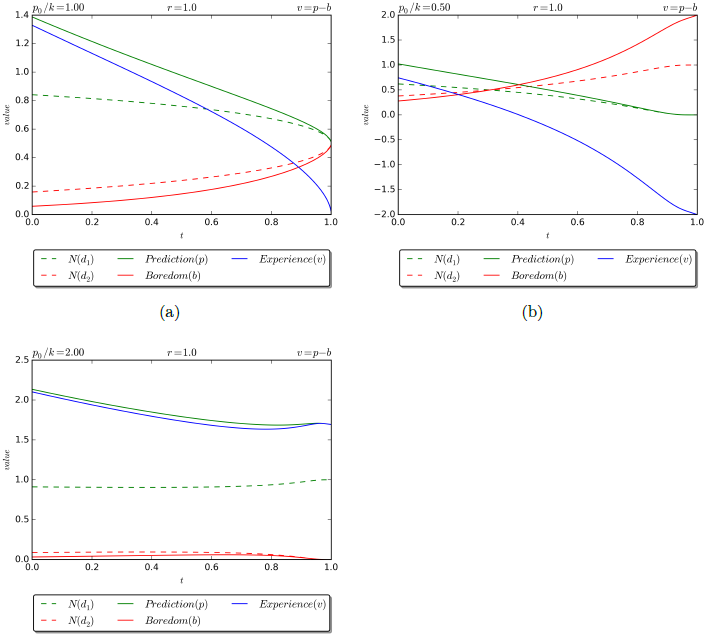
The figure shows the evolution of the probabilities of being in prediction mode, *N*(*d*_1_), boredom mode *N*(*d*_2_), the prediction pleasure (*p*), the boredom-related pain (*b*) and the experience value *v* for a world with high normalized complexity 
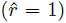
 In Figure 2-*a* there is no initial bias, (*p*_0_ = *k*), in Figure 2-*b* there is a bias that favors exploring triggered by boredom (*p*_0_*/k <* 1) and in Figure 2-*c* the bias is versus prediction against boredom (*p*_0_*/k >* 1). Figure 2-*a* shows boredom increasing to reach the same value as prediction error, bringing the experience to 0. The boredom component can be understood as a signal to explore the world, which here has a rich structure. Figure 2-*b* shows that the overall experience is negative because the agent has a bias for exploring and the world is rich in structure and it is therefore worth exploring the world rather than copy it. In Figure 2-*c* the experience remains positive because the agent has a bias to predict. However, the final experience value will be less than the experience value at 0, this is because the world may contain surprises or unpredictable events. Note that this is different from the dark room situation depicted in Figure 1-*c*, where the agent is a copier in a world that is easy to copy and therefore it will increment his experience by just doing that.

## 4 Discussion

In the predictive coding framework the brain tries to infer the causes of the body sensations based on a generative model of the world. This inverse problem is famously formalized by the Bayes rule. The idea behind this model is that somewhere in the brain there is a decision signal that encodes hypothesis about the sensorial information that is being processed. When incoming sensorial data fully agree with beliefs, prediction error signal becomes stationary. Thus, the system reaches an equilibrium characterized by sampling data from the environment in such a way that the system is never surprised.

A recurring critic of predictive coding is that agents that minimize surprise as the free energy principle mandates, could not possibly engage in explorative behavior or creativity. In a recent update of the theory ^7^, the utility function that would explain agent decision making is defined as the relative entropy or Kullback-Leibler divergence between the probability distributions of likely states and desired states (Schwartenbeck et al., 2013). Both distributions are conditional, the former on empirical priors and the last on priors that represent desired states which are fixed and do not depend on sensory input. In this schema agents will always try to visit the desired states in order to minimize the distance between the desired and likely outcomes.

However, the two major limitations of the free energy principle are still standing. First, the problem of arbitrariness in assigning prior probabilities is never considered. Jaynes’ (Jaynes, 1968) principle of maximum entropy was conceived to specifically addressed the subjectivity problem in assigning prior probabilities. In (Schwartenbeck et al., 2013) this principle is used to convey the idea that the agents that minimize surprise can also have explorative behavior. In a situation such that the agent has not preferred states i.e. the distribution of the desired states is flat, the agent would explore new states since the decision making is unconstrained (flat desired states distribution). However, in the free energy principle, the priors are fixed and do not depend on the sensorial information. This is problematic because an agent with a flat distribution of prior desired states will tend to maximize the entropy over outcomes which is a suboptimal strategy of survival in a world containing a big deal of predictable patterns. Second, if the agent favors specific goal-states, for example, prefers dark and narrow habitats versus wide open spaces, explorative behavior will never occur. Thus, free energy minimization can explain exploration (no goal-state is preferred over other states) and exploitation (goal states are preferred over other states) separately. The interplay between exploration and exploitation can be derived from changes on the precision of the prior over outcome states. However, a mechanistic understanding of the coupling between exploration and exploitaion can not be deducted from the Kullback-Leibler Divergence. The model presented here extends and complements predictive coding by explicitly taking into account the priors and more importantly by adjusting the boredom with the interestingness of the world as it dwindles over time (See Appendix for an example on the evolution of boredom).

Let us illustrate this point with an example. A camper is sitting in front of a bonfire in the woods. It is a chilly and windy night. He hears a noise whose source can not recognize. The camper has two hypothesis to explain the noise, i) the noise is just the breeze moving the leaves or ii) the noise is caused by a Grizzly bear approaching the camp. Let A be the breeze signal and B the bear signal. Initially, since there are only a few bears in those woods and it is a particularly windy night, the camper gives more weight to the hypothesis A -the noise is caused by the wind-than to hypothesis B -it is a bear. Furthermore, the camper enjoys life in general and has a preference to avoid dangerous situations that could put his life at risk. The course of action-stay or go - is given by the divergence between the likely outcomes (the noise is caused by the breeze) and the desired outcomes (it is preferable to be caress by the breeze than eaten by a Grizzly bear).

But let us imagine now that after a long uneventful period of time and the consequent boredom, the camper would like to take the risk of getting into the woods to explore the surrounding area. How can surprisal minimization explains this new behavior? It would need to be possible to readjust the priors (goal-states) in such a way that the agent responds differently to the same stimulus, for example, leaving base camp to explore, rather than staying as the minimization of surprise mandates. More importantly, when the camper decides to explore the woods after being consistently good at predicting the sensory input, it does so because the pleasure of prediction is being overweight by the pain of boredom, resulting in a negative subjective experience that needs to be rebalanced by seeking new states that may bring boredom to lower levels. Crucially, the exhaustion of prediction disrupts the homeostatic balance, which can be counteract by boredom which leads to variety seeking to restore the homeostatic balance. This idea exists in popular parlance in the idiom "die of success", minimizing prediction error would make the organism to seek for easily predictable environments, neglecting exploration and over valuing risk, which would hinder the system’s capacity to prosper and survive in informational complex environments.

From an evolutionary perspective, subjective experience exists to facilitate the learning of conditions responsible for homeostatic imbalances and of their corrective responses. There is an evolutionary advantage in doing surprising actions. For example, in a prey-predator game, both the prey and the predator will have a better change to succeed if they behave surprisingly rather than in predictable ways. Furthermore, if agents always react in the same way to common stimuli e.g. staying if the noise is caused by the breeze, life will be boring and there would be no incentive to explore and discover.

The homeostatic control mechanism that keeps the organism’s internal conditions within admissible bounds reflects the interplay between pleasure associated with prediction and boredom-related pain. Biological systems do not just minimize free energy, rather free energy or surprise is one dependent variable, the other is boredom, and the interplay between both pleasure (prediction) and pain (boredom) defines the independent variable, subjective experience, which is the quantity that systems, all things being equal, maximize.

The importance of boredom needs still to be recognized by researchers. Boredom signals the mismanagement of scarce resources and therefore a better understanding of boredom will have a major impact in economics and behavioral science. We are only just starting to understand the physiological signatures of boredom. Boredom compared with sadness shows rising heart rate, decreased skin conductance level, and increased cortisol levels (Merrifield and Danckert, 2014). Boring environments can generate stress, impulsivity, lowered levels of positive affect and risky behavior. Furthermore, in people with addiction, episodes of boredom are one of the most common predictors of relapse or risky behavior (Blaszczynski et al., 1990). A recent study with humans have shown that a statistically significant number of individuals prefer to administer electric shocks to themselves instead being left alone in an empty room with nothing to do but to think (Wilson et al., 2014). In our model, boredom has a negative effect in the value of the subjective experience, which acts as a catalyzer to explore new states, preventing the organism from "dying of success" by visiting the most likely states in a self fulfilling loop.

The mathematical model here defined conveys the idea that boredom begets creativity. The quantity that organisms maximized is the difference between prediction pleasure and boredom-related pain, and it is through the interplay pleasure and pain, how homeostatic balances and their corrective responses can be acquired and exploited.

We now discuss discuss the implications of our model in relation with reward and valuation systems in the brain. Mobile organisms operate in limited resources environments and therefore need to perform economic evaluations to asses the costs and payoffs of their decisions. Living systems depends for survival on their ability to extract resources to compensate for continuous diffusion of those same resources (Chen, 2012). How they value their internal states (trigger by external cues) to choose a proper course of action is not entirely known.

The midbrain dopaminergic system encode errors in reward predictions. Activity changes in the ventral tegmental area and the substantia negra encode prediction error, more precisely, changes in spike rate encode an ongoing difference between the experience reward and the long term predicted reward (Schultz et al., 1997), (Kakade and Dayan, 2002). In particular, an increase in spike rate represents better than predicted outcomes and decrease denotes worse than predicted (McClure et al., 2003). Thus, activity change in dopaminergic neurons conveys information about prediction error in future reward. The computational model that captures this idea is the temporal differences (TD) learning (Sutton and Barto, 1998),(Markram et al., 2012). That the dopamine system implements TD learning (time discounted sum of expected rewards that can be earned in the future) is uncontroversial.

Montague and colleagues (Montague and Berns, 2002) have suggested that there may exist a generalized valuation function (currency) that goes beyond the expectation violation predicated in TD learning. The orbital frontal stratium would be in charge of providing a common valuation scale or currency. This is the Predictor-Valuation model which is an extension of the reward prediction-error model. Its mathematical formulation is analogous to the Black-Scholes-Merton model including both diffusion and discounting processes. The Predictor-Valuation model, in essence, allows to quantify the expected opportunity costs of being in one state rather than another and preparing one motor action rather than another.

Crucially, opportunity costs require an underlying utility function (Kincaid and Ross, 2009). Our model provides a valuation of the current state discounted by the uncertainty of the environment and the passing of time, that is, taking into account the future states that the agent may visit. The question of which system e.g. a group of neurons in the frontal cortex, dopaminergic pathways, the entire brain, the body etc., we should associate with this utility function is an important question for which we or anybody may have a definitive answer. For an insightful discussion on the conceptual implications of attributing a utility function to a reward system, see (Ross, 2009).

The important point to retain is that although the neurochemistry and neurophysiology of the brain constrain the way in which the reward system is implemented, other aspects can only be explained by reference external to, for example, ecological generalizations about the environment in which the organism evolves and develops. This is taken into account in our model with the discount rate. The model explains why a system that is prone to get bored as much as K, in a world with an informational complexity r will reach its final state. The model is, however, agnostic about the specific policy that the system should carry on. What our model does is to fit the function v of the overall subjective experience (Equation 11). Crucially, if the true optimal could be estimated by the system, then it could use this estimate to update its internal state. This gives the system a way to simulate possible future actions with an overall positive value.

## Footnotes

1 A signal is stationary when its defining probabilities are fixed in time. A signal is ergodic when can be constructed as a generalization of the law of large numbers (long term averages can be closely approximated by averages across the probability space).

2 See the S1 Appendix for the technical definition of surprisal and implications within the free energy principle and the predictive coding framework.

3 Note that surprise, (*H*(*S*)), here refers to the entropy of the sensory states. This is not the same as surprisal, (*H*(*S*|*m*)), in which the probability of the observation *S* is conditional on a model *m*. For a technical discussion of the term surprisal see the appendix and the references wherein.

4 Note that prediction pleasure is the inverse of prediction error, therefore maximize prediction pleasure is the same as minimize prediction error.

5 A utility function is a mathematical description of subjective value that is constructed from choices under incomplete information conditions.

6 *discp*_*T*_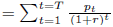, where *discp*_*T*_ is the present value of cash flow in year *T*, *p*_*t*_ is the cash flow, *r* the discount rate and *T* the number of years in the future

7 In truth, the relative entropy or KL distance between the recognition distribution and the generative distribution is included in the seminal paper by Dayan, Hinton and Zemel of the Helmoltz machine (Dayan et al., 1995) which is also used by Friston and collaborators in the free energy principle. The Kullback-Leibler divergence which is always non negative is an upper bound of the quantity that needs to be minimized in the model, namely, the free energy.

## Acknowledgements

This work was possible thanks to Teach Mob Program for visiting professors at the University of Torino. The first author was supported by this program funded the Fondazione Cassa di Risparmio di Torino.

## S1 Appendix

The S1 Appendix file contains the technical definition of Surprisal, a brief introduction to the Free Energy minimization principle and its relevance for predictive coding, discusses the Black-Scholes-Merton equation and provides a demonstration that the variable p (prediction pleasure) in the main text follows a lognormal distribution.

